# Senolytics enhance longevity in *Caenorhabditis elegans* by altering betaine metabolism

**DOI:** 10.1101/2023.12.19.572398

**Authors:** Wenning Lan, Xiaolian Xiao, Xiaojing Zhang, Jingjing Nian, Ziran Wang, Yajiao Wu, Dongcheng Zhang, Junkun Chen, Wenqiang Bao, Chutao Li, An Zhu, Yun Zhang, Fangrong Zhang

**Author notes:** These authors have contributed equally (Wenning Lan, Xiaolian Xiao). **Correspondence should be addressed to:** Dr. An Zhu, Fujian Medical University, Fuzhou, China.; Dr. Yun Zhang, Chinese Academy of Sciences, Xiamen, China.; Dr. Fangrong Zhang, Fujian Medical University, Fuzhou, China.

## Abstract

Aging triggers physiological changes in organisms, which are tightly interlinked to metabolic changes. Senolytics are being developed. However, metabolic responses to natural senescence and the molecular intricacies of how senolytics confer antiaging benefits remain enigmatic. We performed a metabolomics study on natural senescence based on the *C.elegans* model. The results suggest that age-dependent metabolic changes of natural aging occur in *C. elegans*. Betaine was identified as a crucial metabolite in the natural aging process. To explore the common pathway coregulated by different senolytics prolonging nematodes’ lifespan, we fed nematodes three antiaging drugs metformin, quercetin, and minocycline. Our data show that the coregulated metabolic pathways associated with aging include the forkhead box transcription factor (FoxO), p38-mitogen-activated protein kinase (MAPK) and the target of rapamycin (mTOR) signaling pathway, etc. Three antiaging drugs raised betaine levels, consistent with high betaine levels in the younger nematode. Supplement of betaine prolonged the lifespan of nematodes via stimulating autophagy and improving antioxidant capacity. Altogether, our data support proof-of-concept evidence that betaine at appropriate concentrations can extend the lifespan of nematodes.

## INTRODUCTION

The intricate pathophysiology of aging presents a formidable challenge to mitigate age-related mortality and morbidity [1]. The aging process is associated with physiological and molecular degeneration, including the reduction of tissue elasticity and decline in the immune system, anti-infection, and maintaining metabolic homeostasis [2]. In the past decade, the association of host metabolism with the development of aging has been extensively explored. Evidence from studies has revealed metabolic dysbiosis in aging, possibly contributing to aging development [3]. Although solid proof of a causal link between metabolic dysbiosis and aging is still missing, it has been reported that therapeutic strategies such as exercise, diet control and senolytics treatment may confer antiaging benefits through the alteration of metabolic profile [4–6].

Senolytics, a class of drugs designed to eliminate senescent cells, hold promise for promoting antiaging. Drugs such as minocycline, metformin, rapamycin, aspirin and quercetin, were initially prescribed for treating other diseases via modulating host metabolism [7–11], have recently been reported to be effective interventions for counteracting the aging process. Minocycline, a widely used tetracycline analogue for treating acne vulgaris, was reported to have excellent functions of anti-inflammation and attenuate aggregation of proteins [12]. Metformin, a drug extensively used in type 2 diabetes (T2D), has been suggested to increase lifespan and delay age-related diseases [13] through TORC1 inhibition and anti-inflammatory influence on NF-κB. In addition, quercetin is associated with glucose and lipid metabolism in organisms, which has been suggested to regulate the p38-mitogen-activated protein kinase (MAPK) pathway and the insulin-like signaling (ILS) pathway to extend the life span of wild-type nematodes and inhibit NF-κB in turn to reduce the occurrence of inflammation [14]. However, the lack of a comprehensive characterization of natural aging through omics techniques and the limited efficacy of existing senolytic targets hinder their practical clinical application. Thus, how the host metabolism responds to natural senescence and how the metabolic alterations are related to the antiaging benefits of senolytics are urgent to be investigated.

Therefore, this study aimed to detect the metabolic profile alterations on natural senescence based on the *C.elegans* model. We also aimed to pinpoint potential biomarkers associated with the aging process. Additionally, this study explores the potential mechanisms of the beneficial effects of enhancing longevity in nematodes mediated by senolytics, including minocycline, metformin, and quercetin, intending to uncover specific metabolic targets that could potentially be harnessed to slow down the aging process.

## MATERIALS AND METHODS

### Chemical reagents

Dimethyl sulfoxide (DMSO, MACKLIN, AR), 1,1-Dimethylbiguanide hydrochloride (Metformin, Sangon Biotech, purity ≥97.0%), Quercetin (Medchemexpress, purity=98.02%), Minocycline (Medchemexpress, purity= 99.79%)

### Participants

Fasting sera were collected from 100 healthy individuals (male/female, 57/43) with ranged from age 23 to 77 years. Exclusion criteria for the participants were body mass index (BMI) lower than 18.5. This study was approved by the Ethics Committee of Fujian Medical University (ethical code: FJMU2021_181, dtd 12/21/2021). Written informed consent was obtained from all individuals.

### Culture conditions and strains

Wild-type N2 strain worms originally from the Caenorhabditis Genetics Center (CGC, Minneapolis, MN, USA) were used in the present study. They were cultured on Nematode Growth Medium (NGM) agar plates fed with *Escherichia coli* (*E. coli*) OP50 at 20 °C as previously described [15]. Adult hermaphrodite nematodes were collected from the plates into 15 ml tubes, lysed by a fresh bleaching mixture (0.5 mol/L NaOH, 2% NaClO) and washed two times with K medium (32 mM KCl and 51 mM NaCl) (Williams and Chemistry, 2010). Synchronous populations of L1 larval nematodes were collected for further use.

### Drug exposure

Exposure of nematodes to 50 mM metformin, 100 μM quercetin, 100μM minocycline, and single betaine (10 μM, 100 μM, 500 μM, 1mM, 2mM, 10mM, 50mM) was performed from young adults (day 1) until the days of assay endpoints in 6 cm dishes at 20 °C with food of live OP50, 10000 worms per dish. All groups were analyzed in five independent experiments in triplicate. During exposure, nematodes were living in sterile S medium (44 mM KH_2_PO_4_, 100 mM NaCl, 5.74 mM K_2_HPO_4_, 3 μM MgSO_4_, 10 μM potassium citrate, 3 μM CaCl_2_, some trace metals, and 1.29 μM cholesterol). Test solutions were prepared by dissolving different amounts of antiaging drugs in DMSO, with a final working concentration of DMSO of no more than 1%.

### Intestinal lipofuscin levels

After continuous exposure to the drug for 4 or 10 days after synchronization, nematodes were placed in 2% agar pads and anesthetized by levamisole to visualize intestinal autofluorescence. To obtain the relative intensity of the fluorescence from different nematodes, images were collected using a 525 nm bandpass filter at a constant exposure time. Images were taken with an Axio Imager M2 microscope (Zeiss, Oberkochen, Germany) under 100× magnification. ZEN 2 pro software was performed to measure the lipofuscin levels. This assay was performed in triplicate, and 20 nematodes were necessary per treatment.

### Lifespan assay

Synchronized L4 nematodes were exposed to different antiaging drugs in 96-well plates with the liquid S-medium culture system. Live, missing, and dead nematodes were recorded daily, and dead worms were picked out. Survivorship curves were then generated to calculate the percentages of live nematodes in the groups over time. Twenty worms per group were identified in triplicate.

### Locomotion behavior assay

The movement speed of the nematodes was recorded to assess locomotion behavior. After a nematode was slightly touched with platinum wire, a fast movement was defined as when it could move in a sinusoidal motion model continuously in 30 s; otherwise, it was defined as slow movement. After drug exposure in S-medium, transferred nematodes into a new plate without food, and then added 100 μL of M9 buffer to the agar. In a minute, the movement speed was recorded. Twenty nematodes were measured in each treatment, and assays were conducted in triplicate.

### RNA extraction and Real-time quantitative PCR

Worms treated with drugs were collected with M9. Total mRNA was extracted with TRIzol reagent. Amplified cDNA was prepared from approximately 1000 ng of RNA using HiScript II Q RT SuperMix for qPCR (Vazyme, Nanjing, China). SYBR Green Real-time PCR experiments were performed using AriaMx Real-Time PCR (Agilent, California, America) and Taq Pro Universal SYBR qPCR Master Mix (Vazyme). Relative gene expression was normalized to tba-1 mRNA levels. The comparative ΔΔCt method was used to calculate the fold changes in gene expression. The list of primers is provided in Supplementary Table S4. Each experiment was repeated three times, and biological replicate samples were used.

### Observations of autophagy

Worms cultured in drugs were stored in a fixative of 4% glutaraldehyde. After overnight fixation, the nematodes were embedded in agar. Further processing included postfixation in 2% OsO4 and 1.5% potassium ferrocyanide, dehydration in ascending ethanol series followed by acetone and propylene oxide. Samples were embedded in pure epoxy resin 618. Sections were stained with uranyl acetate and examined with a Tecnai Spirit electron microscope (FEI, State of Oregon, America).

### Measurement of ROS

The levels of reactive oxygen species (ROS) in worms were determined by a commercial kit (Beyotime Biotechnology, Shanghai, China). 2’,7’-Dichlorofluorescein-diacetate (DCFH-DA) is effortlessly oxidized to fluorescent dichlorofluorescein (DCF) under the action of intracellular ROS; thus, the levels of ROS were quantified. Accordingly, the worms were cultured in 24-well plates as described above and exposed to 2 mM betaine for 10 days. After the treatment, the worms were incubated with 10 μM DCFH-DA for 2 h at 20 °C without light and washed twice with M9 buffer. Worms were placed in 2% agar pads and anesthetized by levamisole to visualize internal ROS with an Axio Imager M2 microscope (Zeiss, Oberkochen, Germany) and measured by a fluorescence spectrophotometer (λexc = 488 nm, λem = 525 nm). Images were analyzed by ZEN 2 pro software. 20 nematodes were measured in each treatment, and assays were tested in triplicate.

### Assay of antioxidant enzyme activity

After betaine (2 mM) treatment for 10 days, samples were collected. Separately, worms in two groups (DMSO and betaine) were homogenized in cold normal saline for preparing the assay of antioxidant enzyme activity of T-SOD by commercial kits (Nanjing Jiancheng Bioengineering Institute, Nanjing, China). The xanthine/xanthine oxidase method based on the production of O^2-^ anions was used to assay T-SOD activity by ELISA (Biotek, Vermont, America, λ= 450 nm). The liveness of T-SOD is shown as units per milligram of protein (U/mg protein).

### Metabolite extraction

Until analysis, frozen *C. elegans* samples or serum were stored at -80 °C in liquid nitrogen. Serum sample preparation was conducted as described previously [3]. For the analysis, raw serum data from the previous study was used [16]. Elderly adults (day 10) were selected for analysis, and 7000 worms were used for NMR metabolomics. The worm samples were transferred to a precelly (containing 1.4 mm ceramic spheres, MP Biomedicals LLC, Santa Ana, California (SA), America) to obtain metabolites, and then 600 µL of ice-cold methanol and ddH_2_O (2:1) were added to each tube for homogenization by a Tissuelyser-24 (Shanghai Jingxin Industrial Development, China). Using a centrifuge, 12,000 rpm was applied for 30 minutes at 4 °C to obtain the supernatant and then evaporated to obtain a dry metabolite pellet (Eppendorf Concentrator plus, Hamburg, Germany). For the NMR experiments, samples were redissolved in 500 µL of NMR buffer as described [17].

### Data acquisition

A Bruker Avance III HD 600-MHz NMR spectrometer equipped with a TXI probe head was used for NMR metabolic profiling and analysis. With presaturation, ^1^H ^1^D NMR spectra were acquired using Carr–Purcell–Meiboom–Gill (CPMG) pulse sequences (cpmgpr1d, 128 scans, 73728 points in F1, 12019.230 Hz spectral width, 1024 transients, recycle delay 4 s) . NMR spectral data were processed as previously described [17]. Bruker Topspin version 4.0.2 was used to process the data by multiplying the Free Induction Decay (FID) by an exponential window, performing Fourier transformations, and calculating phase differences. To identify the metabolite tissue, MATLAB2014a and Chenomx NMR suite 8.4 were used. Integrations were used to generate the orthogonal partial least squares discriminant analysis (OPLS-DA), permutation analysis, Metabolite Enrichment Analysis (MSEA), and heatmap using MetaboAnalyst 5.0. Q^2^ is a quality assessment statistic that verifies the statistical significance of the identified differences.

### Statistical analysis

GraphPad Prism was used for the univariate statistical analysis (GraphPad Software, La Jolla, CA, USA). Data are represented as the mean ± standard deviation (SD). R framework was performed in the analysis of RNA-seq, and a two-tailed Student’s t-test was used to evaluate statistical significance. When comparing variables pairwise, *p* values were calculated using a two-tailed Student’s t-test. Metabolites with *p* < 0.05 are shown in each supplementary figure. Statistical differences among multiple groups (one-way ANOVA) are indicated by *p* values of < 0.05 (*), < 0.01 (**), < 0.001 (***), or < 0.0001 (****).

## RESULTS

### Age-dependent metabolic changes of natural aging in *C. elegans*

The dysregulation of metabolism is one of the hallmarks of aging. We utilized ^1^H NMR-based metabolomics to analyze and characterize the metabolic alterations in nematodes during their natural aging process. Multivariate statistical analysis was performed to investigate the changes in metabolic profile as *C. elegans* aged. Principle component analysis (PCA) and partial least squares discriminant analysis (PLS-DA) of the nematodes exhibited that the metabolic fingerprints of the nematodes changed with rising age (Figure 1a). When comparing the differences in the metabolic fingerprints between the nematodes with different ages, orthogonal-partial least squares-discriminant analysis (OPLS-DA) revealed increasing correlation coefficients Q^2^ of 0.971 (*p* = 0.01) and an R^2^Y up to 0.998 (*p* = 0.01) (Figure 1b, S1a, and Table S1) with increased clustering. Due to the stochastic nature of nematode aging, notably the onset of mortality after day 10, we excluded data points beyond this timeframe. Nematodes have an approximate lifespan of three weeks, with numerous aging-related characteristics, such as decreased pharyngeal pumping rates, mitochondrial fission, and muscle degeneration, becoming pronounced by day 10 [18]. The heatmap provided a metabolic overview of nematodes as they aged (Figure 1c). To accurately identify metabolite alterations while mitigating potential disruptions due to egg laying, we incorporated young adult nematodes (day 4) and older adult nematodes (day 10) as specific time points for our study. Additionally, we introduced oviposition inhibitors during the entirety of the nematode drug exposure.

**Figure 1.**
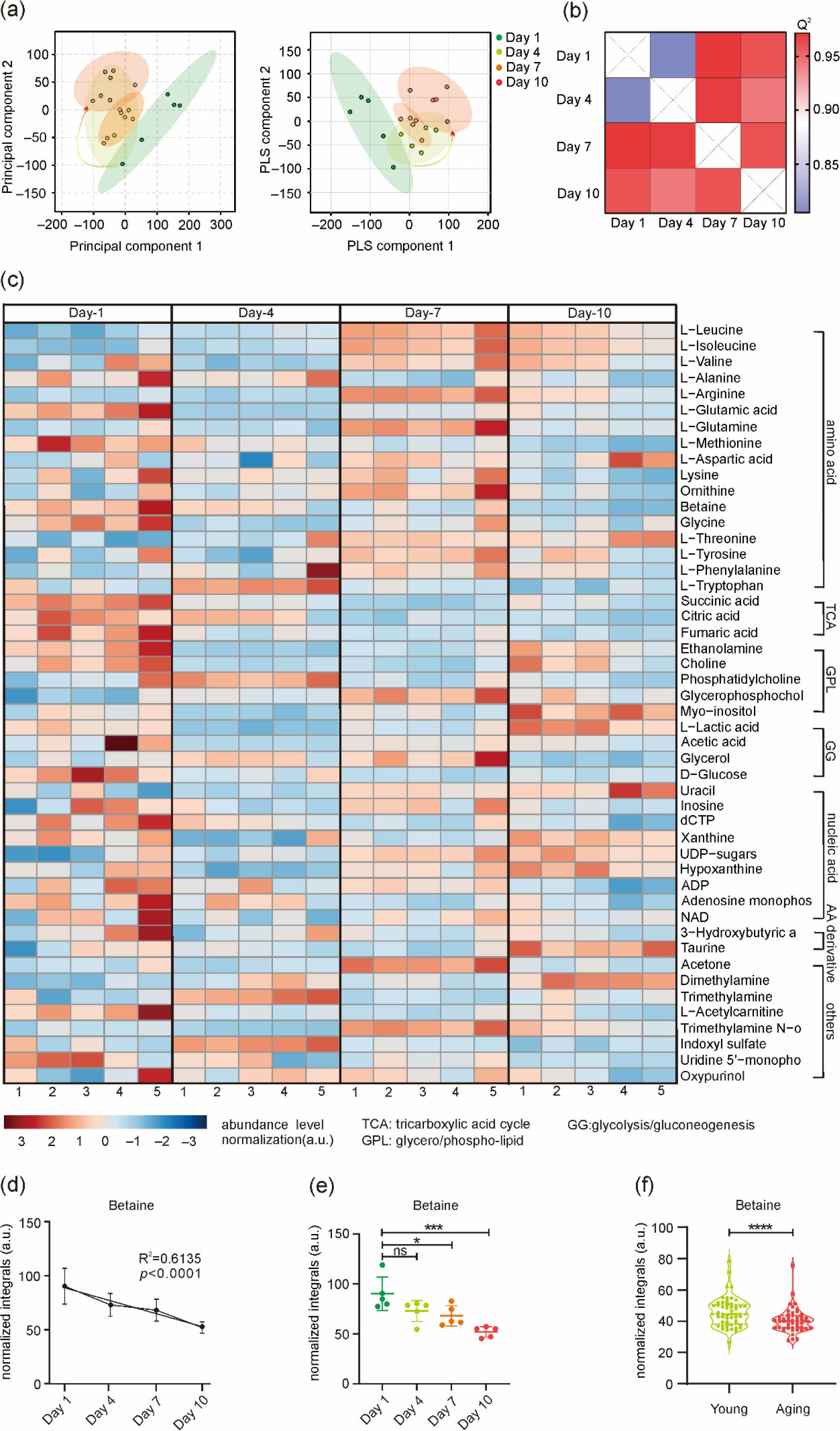
NMR metabolomics analysis of natural aging *C. elegans* samples. (a) PCA and PLS-DA plots of natural aging *C. elegans* samples. The direction of the arrows indicates trends in the metabolism of nematodes from day 1 to day 10. (b) Heatmap exhibiting OPLS-DA-derived Q^2^ for pairwise comparisons of natural aging nematode samples in different ages. (c) Heatmap showing the relative metabolite levels in natural aging *C. elegans* samples by NMR analysis. Each group consists of 1-5 samples. Each column displayed one sample, while rows represented relative metabolites. Red and blue represented increases and decreases. Different classes of metabolites are classified according to their chemical composition and biomolecular. (d) The linear variation of betaine concentration in worms under natural aging. (e) Betaine concentration in nematodes under natural aging. (f) The betaine concentration in human serum samples (60 years is the age limit). * *p* < 0.05, ** *p* < 0.01, *** *p* < 0.001, or **** *p* < 0.001 is significantly different from the control group, and ns shows no significance.

There was a clear distinction between young adults (day 4) and elderly adults (day 10) in the natural aging nematodes, which could be observed in the OPLS-DA plot (Figure S2a). The correlation coefficients provided by the permutation test showed a positive Q² of 0.985 (*p* = 0.02) and R²Y up to 0.985 (*p* = 0.02) (Figure S2b). The reduced NMR spectra analysis displayed specific changes in metabolites, including increases in leucine, isoleucine, valine, lactic acid, arginine, acetic acid, glutamic acid, dimethylamine, ethanolamine, choline, trimethylamine N-oxide, taurine, myo-inositol, uracil, xanthine, uridine diphosphate-sugars (UDP-sugars), and hypoxanthine and decreases in alanine, citric acid, methionine, trimethylamine, lysine, phosphatidylcholine (PC), betaine, glycerol, deoxycytidine triphosphate (dCTP), tryptophan, indoxyl sulfate, and adenosine diphosphate (ADP) (Figure S2c, d). We conducted functional analyses to investigate the age-related alterations in molecular mechanisms. RNA sequencing (RNA-seq) data was analyzed, and the top 20 enriched pathways were exhibited (Figure 2e, Figure S3). Among them, several of the pathways are longevity-related, such as MAPK signaling pathway and longevity regulating pathway, which have been proposed to be related to lifespan [19,20]. At the same time, we identified critical metabolic pathways, such as autophagy, the forkhead box transcription factor (FoxO) signaling pathway, and the target of rapamycin (mTOR) signaling pathway, which are closely related to physiological activities, including autophagy and oxidative stress [21,22] (Figure S3a, S3b). To further identify the metabolites related to aging, correlation analysis was conducted. The positive and negative correlation coefficients represent the metabolite levels increased and decreased with age. Among them, methionine (R^2^ = 0.7533 (*p* < 0.0001)) and betaine (R^2^ = 0.6135 (*p* < 0.0001)) showed the most significant correlation with age (Figure 1d, Figure S3c). Methionine restriction is known to enhance longevity in several model organisms of aging [23,24]. The synthesis of methionine requires a methyl donor. Betaine, as the donor of osmolyte and methyl, is cytoprotective and beneficial to human health [25], which arouses our attention. Our hypothesis suggested that changes in methionine levels could be attributed to the influence of betaine, a compound known to affect an organism’s self-repair mechanisms. Our observations revealed a notable decrease in betaine levels as age increased, and a similar pattern was observed in human samples (Figure 1e and f). The concentration of betaine in the elderly was much lower than that in the young (Figure 1f). Our results implied that betaine might be a potential biomarker for aging.

**Figure 2.**
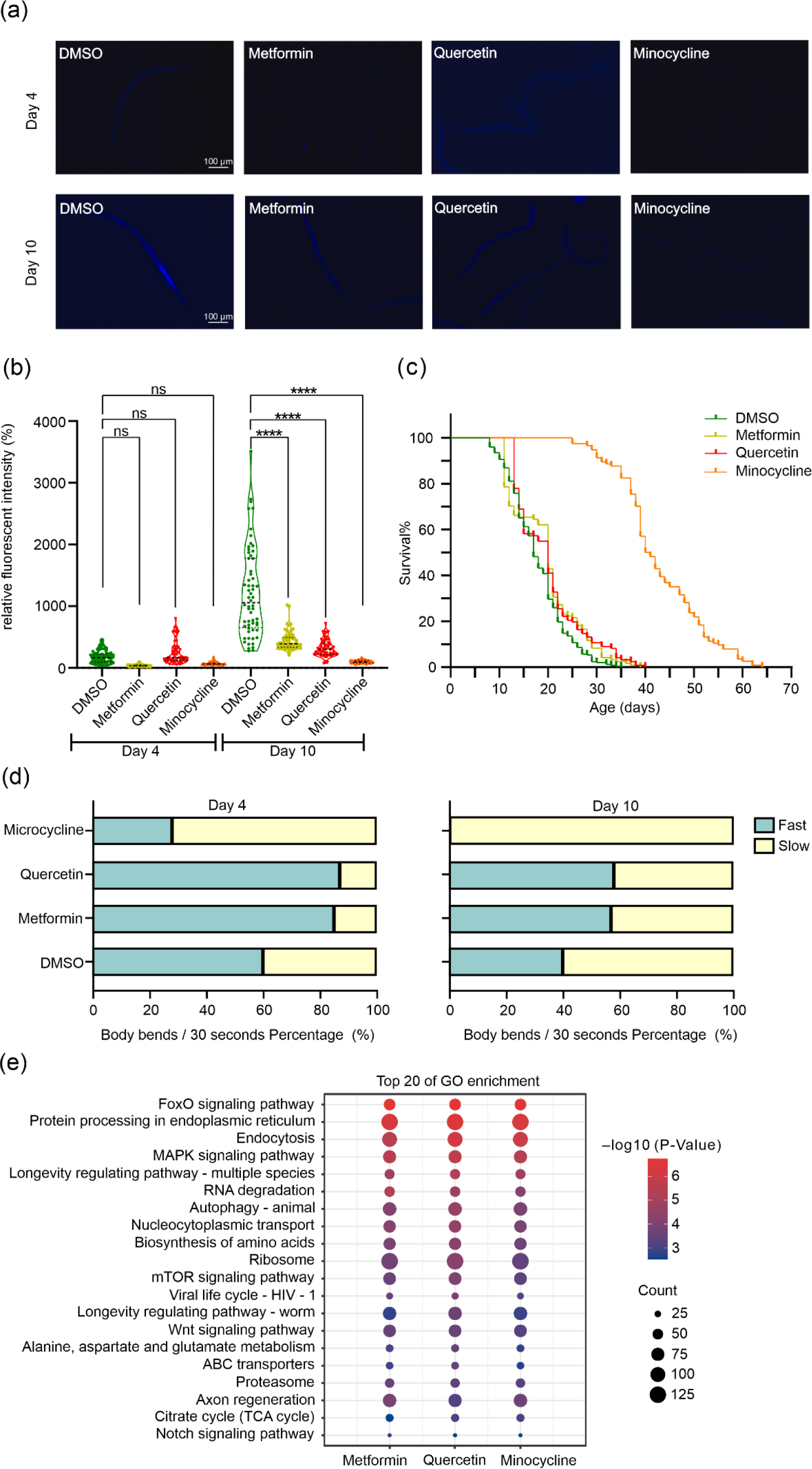
Anti-aging compounds showed lifespan extension. (a) Intestinal fluorescence accumulation in young adults (day 4) and elderly adults (day 10) nematodes treated with DMSO, metformin, minocycline, and quercetin, respectively. (b) Quantitation of the intestinal autofluorescence intensity in young adults (day 4) and elderly adults (day 10) nematodes treated with DMSO, metformin, minocycline, and quercetin. (c) The lifespan of nematodes under the treatment of anti-aging drugs. Median lifespans in this experiment were: DMSO, 35 d; Metformin, 39 d (*p* = 0.003); Quercetin, 40 d (*p* = 0.001); Minocycline, 64 d (*p* < 0.001). (d) Effects on body bends of young adults (day 4) and elderly adults (day 10) nematodes exposed to anti-aging compounds. (e) The Top 20 signaling pathways of nematodes treated with anti-aging drugs in KEGG pathway analysis.

### Effects of senolytics on nematode lifespan extension

To characterize the effects of minocycline, metformin and quercetin, nematodes were divided into four groups and cultured with DMSO, minocycline, metformin and quercetin, respectively. Firstly, to identify the effects of the antiaging drugs on aging delay, an aging-related indicator of lipofuscin accumulation was evaluated. Compared with the DMSO control group (day 10), the relative fluorescence intensity in nematodes with exposure to minocycline, metformin and quercetin (day 10) decreased to 0.08 (*p* < 0.0001), 0.31 (*p* < 0.0001) and 0.27 (*p* < 0.0001) fold, respectively (Figure 2a, b). The results suggested that the antiaging drugs slowed the deposition of metabolic waste products and lipid oxidation in nematodes, and minocycline was the most effective. We further measured the survival of the worms in the four groups. Under the antiaging drugs’ conditions, nematodes’ lifespan was extended to different levels. Compared with the control group, the lifespan of minocycline-fed nematodes was increased by 140.5% (*p* < 0.0001), while metformin and quercetin slightly improved the lifespan by 12% (*p* < 0.001) and 9% (*p* < 0.003), respectively (Figure 2c, Table S2). The results showed that antiaging compounds including minocycline, metformin, and quercetin could extend organisms’ lifespan, consistent with early studies [26]. A distinct feature of aging nematodes is decreased mobility and muscle function. Therefore, we assessed the alterations in the locomotion capacity of worms under the treatment of antiaging drugs. The frequencies of body bending were measured to evaluate the locomotion behavior of nematodes. Our results showed that the frequencies of body bending were significantly increased by 20% in metformin- and quercetin-treated nematodes compared to the control, while slowed in minocycline-fed nematodes (Figure 2d).

Moreover, we obtained the signaling pathways strongly related to the above drugs utilizing gene ontology (GO) enrichment and KEGG analysis. RNA sequencing (RNA-seq) data was analyzed, the results presented top 20 enriched pathways, such as the FoxO signaling pathway, MAPK signaling pathway, autophagy, longevity regulating pathway, and mTOR signaling pathway, which were similar to those involved in natural aging process (Figure 2e, Figure S4). In summary, we confirmed the life-prolonging effects of these three antiaging drugs on *C. elegans* and identified the life-prolonging metabolic pathways co-regulated by them.

### Changes in metabolites correlate with therapeutic benefits

We further focused on the regulation of the antiaging drugs on nematode metabolism to explore their underlying mechanism on the extension of lifespan. Firstly, PCA was performed on the ^1^H-NMR data in response to the senolytics intervention. According to the PCA plot (Figure 3a), the tendency of metabolite variations in elderly nematodes treated with antiaging drugs coincided with that detected in young nematode samples, which might occur due to the regulation of antiaging drugs. The results indicated that the antiaging drugs could intervene in the metabolic profile of aged nematodes and alter their metabolome toward a more youthful state. OPLS-DA analysis displayed an evident separation between the metabolomes of samples from elderly controls (day 10) and senolytics groups, indicating that the metabolic profiles of the nematodes were significantly changed compared with the elderly controls (Figure S5a-S5c). The differential metabolites between elderly controls (day 10) and senolytics groups were identified. The intervention of senolytics on the differential metabolites was supplemented in Figure S5d-S5i. Our results showed that 16 differential metabolites were common in three comparisons (minocycline *versus* elderly control, metformin *versus* elderly control and quercetin *versus* elderly control) (Figure S5j-S5l), including citric acid, succinic acid, glycerophosphocholine, betaine, L-alanine, trimethylamine, phosphorylcholine, L-Glutamine, ethanolamine, L-tryptophan. Among them, betaine was prominent, which was also identified as aging-related metabolite in natural aging nematodes in our study. The levels of betaine in the four groups were compared, as figure 3b illustrates, betaine was significantly increased after senolytics administration (*p* < 0.05), and the highest concentration was observed in minocycline-fed nematodes (*p* < 0.0001). Notably, metabolites like phosphorylcholine, ethanolamine and glycine, which were involved in the betaine metabolic pathway, remarkably increased after senolytics administration (Figure S6). These results indicated that the three antiaging drugs could modulate betaine metabolism in aged nematodes, and betaine might be an essential metabolite for the antiaging benefits of senolytics.

**Figure 3.**
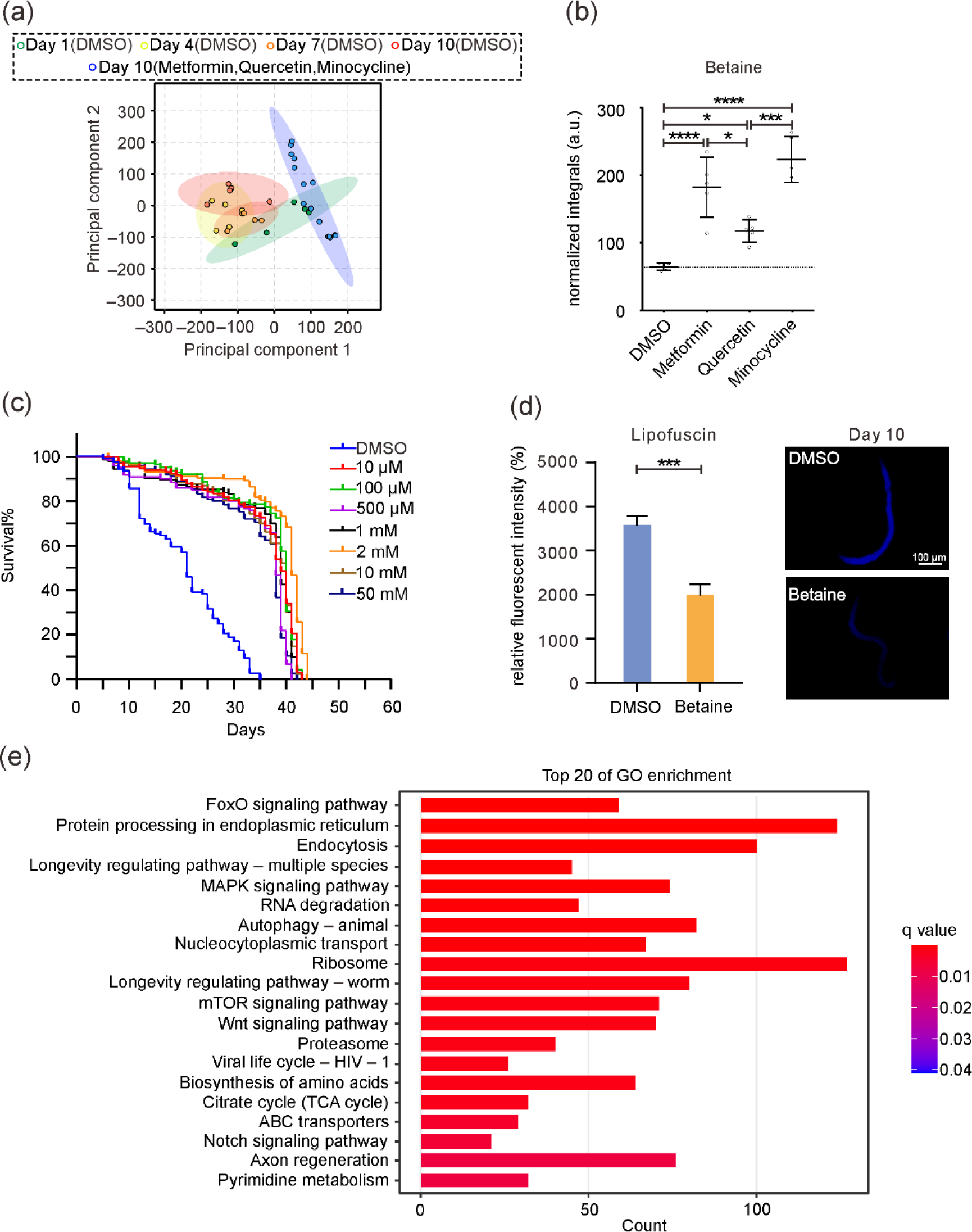
Enhancement of lifespan by betaine in nematodes. (a) PCA plot of nematode samples in natural aging and anti-aging compounds cultured on day 10. (b) Statistical analysis of betaine in nematode samples cultured in anti-aging drugs. (c) The lifespan of nematodes under the treatment of betaine in different concentrations. (d) A representative image of the intestinal cells of nematodes under exposure to betaine shows the accumulation of autofluorescent lipofuscin. Scale bars are 100 µm. (e) The Top 20 signaling pathways of nematodes treated with betaine in KEGG pathway analysis. Count means the number of genes.

To assess the impact of betaine on the lifespan of nematodes, survival analysis was performed by applying different concentrations of betaine. According to the survival curves, betaine in different concentrations exhibited various impacts on extending the lifespan of nematodes compared to the DMSO group. The most significant effect was observed at a concentration of 2 mM betaine treatment (Figure 3c), in which the lifespan of nematodes increased by 114% compared to the controls (*p* < 0.0001, Table S3). The relative fluorescence intensity of nematodes was significantly decreased in the betaine group (2 mM) (Figure 3d), which suggested that betaine had similar effects with antiaging drugs in delaying lipofuscin accumulation. To identify the mechanism underlying the life-prolonging effects of betaine, we analyzed RNA sequencing (RNA-seq) data and obtained the top 20 metabolic pathways with significant changes (Figure 3e, Figure S7). The metabolic pathways regulated by betaine were similar to that held by antiaging drugs, such as the FoxO signaling pathway, MAPK signaling pathway, autophagy, longevity regulating pathway and mTOR signaling pathway, suggesting betaine may play a role in longevity regulation and physiological activities like autophagy and oxidative stress.

### Autophagy is required for the beneficial effects of betaine

Most antiaging interventions depend on autophagy to exert their protective properties, and betaine has been reported to stimulate autophagy [27]. Accumulating evidence underscores the pivotal role of autophagy in influencing lifespan through various vital physiological processes. This evidence suggests that enhanced autophagy has health benefits and can significantly contribute to its function of retarding the aging process [28]. Thus, we investigated whether autophagy plays a role in the beneficial effects of betaine in the lifespan of nematodes. The mRNA expression of several autophagy upstream target genes, such as *aak-2*, *rheb-1*, and *let-363*, were measured by RT-qPCR. Compared with the elderly control group, the mRNA expression of *rheb-1* and *let-363* in nematodes treated with betaine (2 mM) decreased to 2.9 (*p* < 0.0001) and 3.9 (*p* < 0.001) folds, respectively (Figure 4a), while *aak-2* increased to 4.9 fold (*p* < 0.01). The results suggested that the autophagy-related genes regulated *C. elegans* longevity in response to betaine. We further investigated the formation of autophagosomes in the elderly controls and betaine group using electron microscopic analysis. Increased autophagosome under betaine feeding was observed (Figure 4b). Our results indicated that the beneficial effects of betaine might be partially due to the activation of autophagy through the AMPK/mTOR signaling pathway.

**Figure 4.**
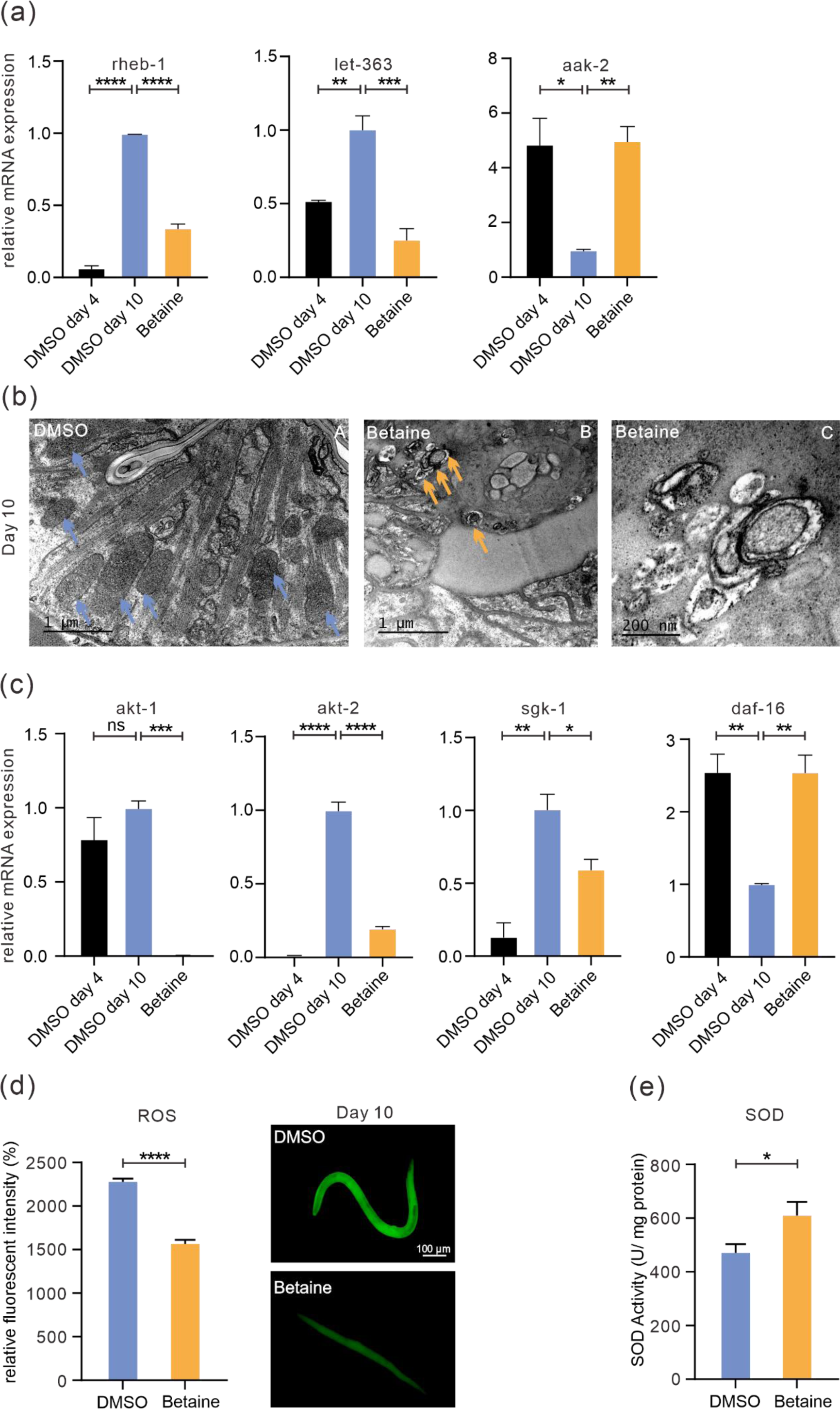
Effects of betaine exposure on autophagy and oxidative stress responses in nematodes. (a) Comparison of the expression pattern of genes demanded autophagy in young (DMSO Day 4), elderly (DMSO Day 10), and betaine-treated (Day 10) nematodes, the outcomes were presented as the relative expression rates between tba-1 reference gene and targeted genes (*rehb-1*, *let-363*, *aak-2*). (b) The structure of autophagy was analyzed by electron microscopy experiment. Transmission electron microscopy of nematodes fed with DMSO (A) and betaine (B), in turn, and higher magnifications of the boxed region of B was shown in C. Blue arrows indicate mitochondria, and yellow arrows indicate autophagosomes. Scale bars are 1 µm in A and B, and 200 nm in C. (c) Comparison of the expression pattern of genes demanded oxidative stress responses in young (DMSO Day 4), elderly (DMSO Day 10), and betaine treated (Day 10) nematodes, the results were presented as the relative expression ratio between tba-1 reference gene and targeted genes (*akt-1*, *akt*-2, *sgk-1*, *daf-16*). (d) The left side shows the relative fluorescence intensity comparison between the control and betaine-exposed groups. The right panels are images of nematodes showing the generation of ROS. The scale bar is 100 μm. (e) The comparison of antioxidant enzyme activities between the control group and betaine exposed group. Data were expressed as the mean ± SEM. of three independent experiments. \**p* < 0.05, \*\**p* < 0.01, \*\*\**p* < 0.001, and \*\*\*\**p* < 0.0001 indicated a significant difference when compared to the elderly (DMSO Day 10), and n.s. was not significant.

### Betaine reduced oxidative stress of *C. elegans*

A key feature of aging is increased oxidative stress, which refers to elevated levels of reactive oxygen species (ROS) [29]. To assess whether oxidative stress is involved in betaine prolonging nematode lifespan, we analyzed the changes in the mRNA expression of oxidative stress related genes (*akt-1*, *akt-2*, *sgk-1* and *daf-16*) after treatment with betaine (2 mM). After administering betaine treatment, the mRNA expression of these genes showed notable alterations, shifting toward a resemblance with the young group rather than the elderly group, as opposed to the control group (Figure 4c, *p* < 0.05). Betaine is trimethylglycine with antioxidant properties, and it has been reported to attenuate oxidative stress in rat models of Alzheimer’s disease [30]. We monitored ROS production in the elderly controls and betaine group using H2DCF-DA. A significant reduction of ROS accumulation in nematodes treated with betaine was observed, whose dichlorofluorescein (DCF) fluorescence intensity was 30% lower than that in the elderly controls (*p* < 0.0001) (Figure 4d). The antioxidant enzyme superoxide dismutase (SOD), is directly linked to intracellular defence against reactive oxygen species [31]. We examined the effect of betaine on SOD activities. The betaine group exhibited a higher SOD activity, which comes to 1.3 fold that of the control (*p* < 0.05) (Figure 4e). These results indicated that betaine could inhibit ROS accumulation in *C. elegans*. The free radical theory of aging points out that the formation of ROS and the subsequent proliferation of damaged macromolecules mainly account for aging [32]. Therefore, we proposed that betaine retard the aging process may be partially due to its ability to scavenge ROS. Overall, the prolongation of the nematode lifespan by betaine might be closely related to its antioxidant activity, which correspondingly improved the resistance of *C. elegans* to oxidative stress.

## DISCUSSION

Evidence shows that aging influences various biological pathways, including metabolism, and leads to organ failure and even death. For aging assessment, the decline in memory and cognition, hypertrophic growth in cardiomyocytes, and facial recognition are commonly used [33]. However, the above methods are easily affected by the external environment and individual differences and cannot be used as universal detection methods. Identifying generalizable means of assessing aging is crucial to supply information on the aging process, as they illustrate the state of aging in organisms and provide a method for extending lifespan. Untargeted NMR metabolomics, a powerful tool with simple interpretation and high reproducibility, helped us identify the related metabolites in aging.

During aging, the metabolite levels of nematodes showed significant changes. In our results, amino acids, such as methionine, alanine, betaine and tryptophan, decreased in the elderly compared to young samples. In contrast, glycerophospholipids like choline and ethanolamine were increased in the elderly. Interestingly, we noticed that methionine, choline and betaine act as methyl donors and have been proposed to modify age-associated DNA methylation and subsequently alter the age-associated physiologic and pathologic processes [34]. A clear negative correlation between tryptophan levels and age was evident, aligning with the observed trend in human serum samples [35]. Previous research has reported that tryptophan levels were often lower in aging-related disorders [36–38], consistent with our results. Strikingly, metabolites related to the tricarboxylic acid (TCA) cycle were also altered in nematode samples of different ages. The diminishing demand for succinic acid, citric acid, and fumaric acid in aging is quite evident as nematodes age, possibly due to the mitochondrial components being impaired by mitochondrial ROS with aging [39].

In this study, we found that betaine, a key metabolite, not only serves as a common metabolic marker of natural aging but also belongs to a crucial metabolic pathway coregulated by antiaging drugs, essential in nematodes’ lifespan extension. Our study showed that betaine might be an essential metabolite for the senolytics treatment’s antiaging benefits, and several metabolites involved in the betaine metabolic pathway also exhibited significant alterations in response to the drugs. Therefore, we built a biological model for the betaine metabolic pathway to explain the potential mechanism (Figure 5). For one critical branch of this pathway, betaine is transformed from choline, whose synthesis benefits from some important compounds, including ethanolamine, PE and PC. We focused on this branch because choline, ethanolamine, PE and PC are essential chemicals involved in aging delaying [40]. Ethanolamine (Etn), a precursor of phosphatidylethanolamine (PE), was found to be upregulated in the antiaging process, which further led to the increase of phosphatidylcholine (PC) (Figure S6). However, stable choline concentrations in nematodes cultured in antiaging drugs were observed, possibly due to conversion to betaine (Figure S6). In early studies, PE is essential for the autophagic process; a higher concentration of PE helps delay aging [41]. PC is obtained from diet, which is the primary source of choline. Supplementation with PC has been verified to help slow the aging process and extend the lifespan of nematodes [40]. Interestingly, glycine, a downstream metabolite of the pathway, also showed an increased trend after antiaging treatment, especially in the group of metformin (p<0.001) (Figure S6), indicating an upregulation of this pathway. Recent studies have revealed that glycine supplementation contributes to extending the life of nematodes [42].

**Figure 5.**
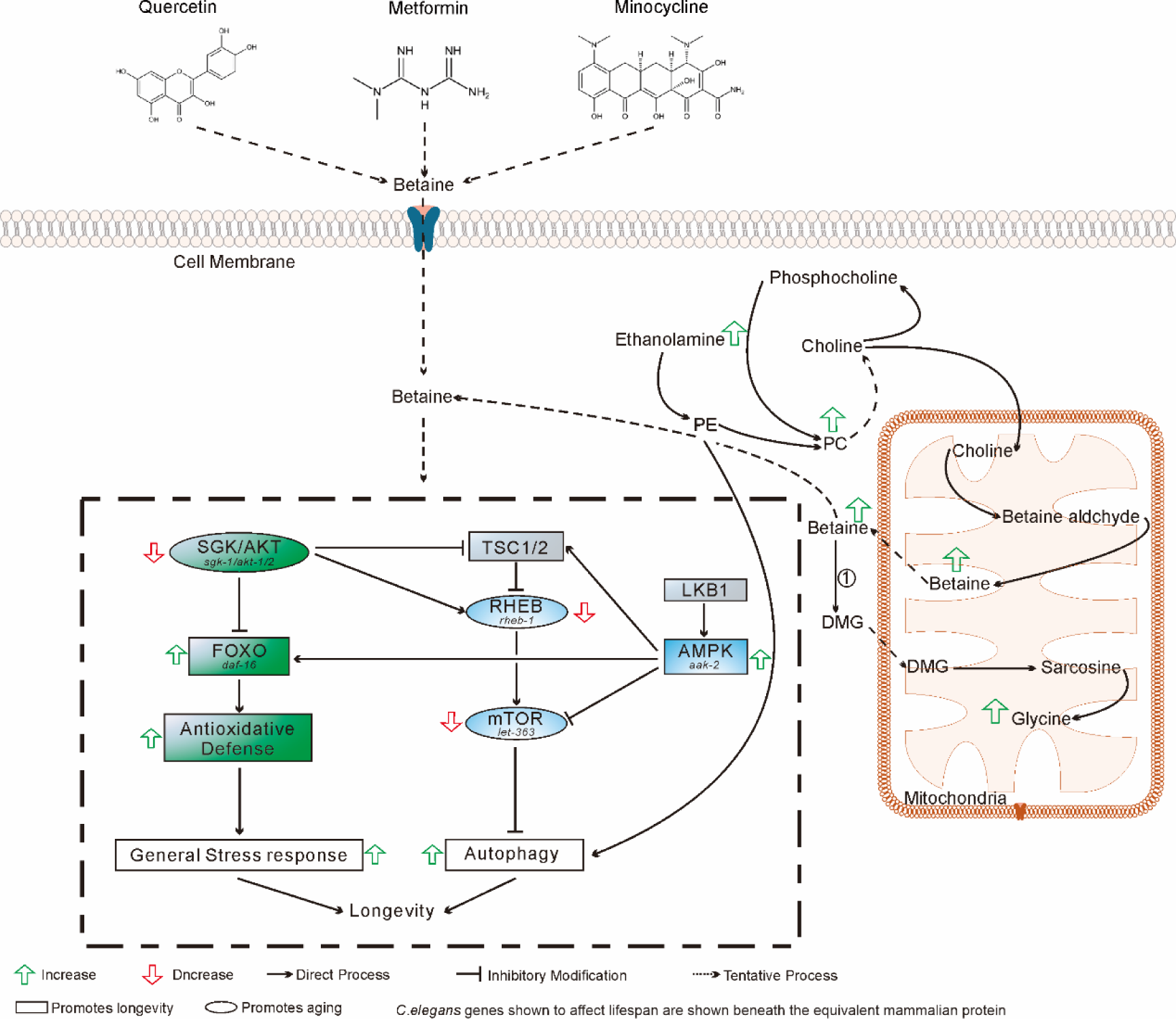
An abridged view of the synthesis and metabolism of betaine and its role in the age-related pathway in *C. elegans*. The enzymes mentioned are displayed and labeled in the cycle with individual numbers: 1. BHMT, Betaine-homocysteine methyltransferase; DMG, N,N-dimethylglycine; PE, Phosphatidylethanolamine; PC, phosphatidylcholine. Genes of nematode that affect lifespan are shown below equivalent mammalian proteins. Proteins not yet exhibited to influence aging are shown in gray.

In our study, betaine stimulated autophagy by inhibiting the mTOR pathway. In early research, autophagy occurred at baseline levels in tissues under normal growth conditions, but it was re-regulated promptly in response to other situations [43]. For instance, betaine-controlled expression levels of *rheb-1*, *let-363*, and *aak-2* were consistent with those in the young group, which played a crucial role in prolonging the lifespan of creatures and has been shown to stimulate autophagy to eliminate senescent cells [44]. Our study also observed these autophagy-related genes altered in response to betaine. In worms, *let-363* encodes an ortholog of mTOR, a highly conserved kinase that regulates autophagy [45]. Studies on *C. elegans* have noticed that suppressed expression of mTOR/*let-363* was associated with an increased lifespan of almost more than double [46]. The *rheb-1* switches to toggle the activity of TOR complex 1 (TORC1) between anabolism and catabolism, thus controlling lifespan, development and autophagy [47]. Honjoh et al. found that a decrease in *rheb-1* prolongs the lifespan of the worms [48]. The *aak-2* encodes the α-subunit of AMPK, which is critical for autophagy, stress responses, energy metabolism and life span [49,50]. Onken B et al. indicated that metformin could increase the expression of *akk-2* and extend lifespan through the AMPK/*akk-2* pathway [51]. In addition, as previously reported, the AMPK/mTOR signaling pathway protects the expected initiation of autophagy and delays the organism’s ageing [22]. However, there is no clear understanding of how autophagy promotes lifespan extension.

In addition, Our study shows that the antioxidant capacity of nematodes was improved under betaine treatment by promoting the FoxO pathway. Oxidative stress related genes, including *akt-1*, *akt-2*, *sgk-1* and *daf-16*, showed significant alterations, shifting toward the young group. The *daf-16* gene encodes the FoxO transcription factor, which has been reported to significantly regulate the aging process in both *C. elegans* and mammals [52]. Liu *et al.* indicated that lentinan could increase the expression of *daf-16* and dramatically prolong the *C. elegans* lifespan [53]. The *akt-1/2* genes encode the kinase cascade in the insulin signalling pathway, which has been identified to be required for the control of longevity [54]. Liu *et al.* observed a decrease in *akt-1* and *akt-2* expressions after paeoniflorin administration, which showed a beneficial effect on lifespan [55]. The *sgk-1* encodes a threonine/serine protein kinase that is crucial in the cellular stress response, and is also thought to act similarly to *akt-1/2* in lifespan control by phosphorylating and inhibiting the nuclear translocation of DAF-16/FoxO [56]. Study on *C. elegans* noticed that suppressed expression of *sgk-1* resulted in increased stress resistance and an extension of lifespan [57].

Despite the metabolic patterns of aging have been gathered, some limitations to this study exist. Our study recruited 5dishes (10000 worms per dish) and the limited sample size might prevent detection of differences with smaller effect sizes. Though a close association was found for betaine and aging, the levels of betaine in various longevity mutants, such as daf-2(-) were lacking, which will be more persuasive to support our hypothesis. Lastly, future studies could also explore whether betaine can be advantageously used in humans.

In conclusion, Based on this study, we identified the metabolic reprogramming of nematodes due to natural aging. In addition, our study revealed that the antiaging drugs significantly affect anti aging by regulating betaine level and related target genes. Moreover, the beneficial regulation of betaine relies on the participation of autophagy and antioxidative activity (Figure 5). Our findings reveal the relationship among senolytics, betaine and aging. At the same time, this study provides a reference for subsequent clinical trials, and hopefully, betaine is a promising intervention against aging.

## Acknowledgements

This work was supported by Startup Fund for High-level Talents of Fujian Medical University (XRCZX2021020), Natural Science Foundation of Fujian Province (2022J01660, 2021J05045) and National Natural Science Foundation of China (82203431, 82104520), Key Research Program of the Chinese Academy of Sciences (ZDRW-CN-2021-3), Fujian Science & Technology Innovation Laboratory for Optoelectronic Information of China (2020ZZ114).

## Author Contributions

**Wenning Lan:** Conceptualization, Methodology, Investigation, Data curation, Formal analysis, Visualization, Writing–original draft, Writing–review & editing. **Xiaolian Xiao:** Investigation, Data curation, Formal analysis, Visualization, Writing–review & editing. **Xiaojing Zhang:** Investigation. **Jingjing Nian:** Investigation. **Ziran Wang:** Visualization. **Yajiao Wu:** Investigation, Formal analysis. **Dongcheng Zhang:** Investigation, Formal analysis. **Junkun Chen:** Investigation, Formal analysis. **Wenqiang Bao:** Methodology**. Chutao Li:** Methodology. **An Zhu:** Conceptualization, Methodology, Data curation, Resources, Supervision, Project administration, Funding acquisition. **Yun Zhang:** Conceptualization, Methodology, Data curation, Resources, Supervision, Project administration, Funding acquisition. **Fangrong Zhang:** Conceptualization, Methodology, Formal analysis, Resources, Data curation, Writing – review & editing, Supervision, Project administration, Funding acquisition.

## Ethical Statements

This study was approved by the Ethics Committee of Fujian Medical University (ethical code: FJMU2021_181, dtd 12/21/2021).

## Data availability

This study includes no data deposited in external repositories.

